# Efficacy of 60 personal mosquito repellents and perfumes against *Aedes aegypti* mosquitoes

**DOI:** 10.64898/2026.08.27.747522

**Authors:** Perran A. Ross, Véronique Paris, Apeksha L. Warusawithana, Kirby D. X. Romualdez, Ashritha Prithiv Sivaji Dorai, Jasmeen Kaur, Mohd Farihan Md Yatim, Luyang Li, Ary A. Hoffmann

## Abstract

Personal insect repellents can offer an effective means of preventing mosquito bites. Despite a wide variety of products that claim to repel mosquitoes, relatively few studies have systematically compared their performance through standardized tests. In Australia, chemical-based repellents must be registered with the Australian Pesticides and Veterinary Medicines Authority (APVMA); however, many unregistered products with unverified effectiveness remain commercially available. Here, we tested 55 insect repellents and 5 non-repellent perfumes and deodorants sold in Australia for their protection against *Aedes aegypti* mosquito landings (as a proxy for bites) using arm-in-cage assays across eight human subjects. We tested 43 topical repellents (containing 18 different primary active ingredients), four wristbands, three clothing stickers, three ultrasonic devices, one garment, one repellent applied to fabric, two perfumes, two deodorants and one antiperspirant. Thirteen repellents, including all wristbands, stickers and ultrasonic devices did not provide complete protection against mosquito landings for any human subject, while a further sixteen provided complete protection for 1 hr or less on average. Repellents containing DEET, oil of lemon eucalyptus, picaridin and IR3535 as the primary active ingredient were the only products that provided more than 2 hours of complete protection, but with significant variation among products with the same active ingredient. Only 7/30 essential oil-based topical repellents and perfumes provided complete protection for greater than 1 hr. Nearly half (25/52) of the repellents subject to registration with the APVMA were unregistered despite being advertised or labelled as mosquito repellents, and these tended to be less effective than registered products. Only 5/28 repellents met or exceeded their claimed protection time. This study highlights the prevalence of unregistered and ineffective mosquito repellents on the Australian market while providing evidence for products and active ingredients that do protect against mosquito bites under laboratory conditions.

## Introduction

The prevalence of mosquito-borne pathogen transmission is increasing globally (George et al., 2024). In Australia, mosquitoes cause significant nuisance biting and transmit pathogens including Ross River virus, Barmah Forest virus and *Mycobacterium ulcerans* (Buruli ulcer) that each cause hundreds of cases of disease annually (Hosen et al., 2026; Mee et al., 2024). Controlling mosquito populations remains the most effective means to reduce mosquito-borne pathogens transmission and nuisance biting at broad scales. Mosquito populations can be suppressed through chemical and biological insecticide applications (van den Berg et al., 2012), aquatic habitat management (Tusting et al., 2013), mass trapping of adults (Barrera, 2022) and community-based interventions (Heintze et al., 2007). In recent years, programs involving the release of modified mosquitoes including sterile and incompatible males (Dobson, 2021), genetically modified strains (Spinner et al., 2022) and mosquitoes transinfected with *Wolbachia* endosymbionts (Moretti et al., 2025) have also been successful in reducing populations and/or pathogen transmission. Household measures to control mosquitoes and prevent bites include the use of screens (Che-Mendoza et al., 2018), bed nets (Curtis et al., 2006) and spatial repellents such as mosquito coils (Chen et al., 2025).

Personal mosquito repellents can be an effective way for individuals to protect themselves against mosquito bites and reduce the risk of contracting a mosquito-borne disease (Frances & Debboun, 2022). Topical repellents containing high concentrations of DEET (Diethyltoluamide), picaridin, or oil of lemon eucalyptus (containing para-Menthane-3,8-diol or PMD) remain the gold standards, with strong evidence for long-term protection against mosquito bites under laboratory and field conditions (Fradin, 1998; Goodyer et al., 2010; Nguyen et al., 2023; Webb & Hess, 2016). Other active ingredients including IR3535 (Ethyl butylacetylaminopropionate), citronella oil and 2-undecanone also show repellent efficacy (Nguyen et al., 2023), but with fewer studies available demonstrating long-term protection. Permethrin-treated clothing can also provide protection against mosquito bites (Banks et al., 2014), though other wearable products including wristbands, stickers and ultrasonic devices have little to no evidence of efficacy despite being widely available (Revay et al., 2013; Rodriguez et al., 2017; Toma et al., 2024; Webb & Russell, 2011).

Products sold in Australia that claim to repel mosquitoes must be registered with the Australian Pesticides and Veterinary Medicines Authority (APVMA) (Webb & Hess, 2016), though non-chemical products including ultrasonic devices are exempt. Registration requires evidence demonstrating the efficacy and safety of the product (https://www.apvma.gov.au/registrations-and-permits). Nevertheless, many unregistered mosquito repellents are sold online through retailers such as Amazon, Etsy, Temu and iHerb and independent retailers. Under Australian consumer law, products are expected to work as claimed (https://www.accc.gov.au/business/selling-products-and-services) but there is a need for independent testing. This is best exemplified by recent trials conducted by choice.com.au revealing that most sunscreens sold in Australia did not meet the advertised SPF standard (https://www.choice.com.au/health-and-body/beauty-and-personal-care/skin-care-and-cosmetics/articles/sunscreen-test). For mosquito repellents, claims of protection time (e.g. up to 8 hours of protection) are often included on the label, but information is lacking on how protection times were derived and whether they are comparable among similar products.

Standardised tests can provide valuable information on the efficacy of mosquito repellents due to their independent nature and use of controlled conditions (Luker, 2024). Many past studies have compared multiple repellent products under the same conditions, highlighting variation in the efficacy of different products and active ingredients (Barnard & Xue, 2004; Chou et al., 1997; Fradin & Day, 2002; Masetti & Maini, 2006; Peng et al., 2022; Rodriguez et al., 2015). While a small number of products sold in Australia have been tested independently (Frances et al., 2009; Frances et al., 2005; Ritchie et al., 2006; Webb & Russell, 2011), there is limited recent information despite many products appearing on the market in the last decade.

Here, we aimed to test a broad range of mosquito repellent products under standardized conditions with a range of active ingredients and modes of application across a consistent set of human subjects. The repellents included both products registered with the APVMA and unregistered products. Additionally, we tested a small range of non-repellent perfumes and deodorants since these can contain compounds that repel mosquitoes (Rodriguez et al., 2015; Zeng et al., 2018). We performed tests with *Aedes aegypti* mosquitoes that are widespread in northern Queensland, Australia using arm-in-cage assays, providing a balance between biological realism (with a human arm used as the attractant) and controlled conditions with a paired design, while being relatively high-throughput (Luker, 2024).

## Materials and methods

### Mosquito repellent products

We conducted a comprehensive search for insect repellent products available in Victoria, Australia in store and online in 2025. We selected products with a broad range of active ingredients, modes of application and product categories, including topical repellents (e.g. sprays, lotions, creams, wipes), wristbands, clothing stickers, ultrasonic devices and clothing. We first searched for products containing key active ingredients known to provide strong protection against mosquito bites, including DEET (Diethyltoluamide) (Leal, 2014), picaridin (Goodyer & Schofield, 2018), oil of lemon eucalyptus (which contains p-Menthane-3,8-diol or PMD)(Carroll & Loye, 2006) and IR3535 (Ethyl butylacetylaminopropionate) (Thavara et al., 2001). We then expanded our search to include products containing naturally derived compounds including essential oils that have also been shown to repel mosquitoes (Drapeau et al., 2009; Luker et al., 2023; Maia & Moore, 2011; Mitra et al., 2019). We also searched for products containing other compounds with demonstrated efficacy against mosquito bites, including delta-undecalactone (Menger et al., 2014), 2-undecanone (Witting-Bissinger et al., 2008), nootkatone (Fernandez Triana et al., 2025), catnip (Melo et al., 2021) and coconut fatty acids (Zhu et al., 2018), but no products containing these compounds were available in Australia at the time of the study. We also searched the APVMA website (https://portal.apvma.gov.au/pubcris) for products that had been registered for use as an insect repellent. Finally, we extended our search to other products that may repel mosquitoes despite not being labelled as mosquito repellents, including perfumes containing hedione (methyl dihydrojasmonate) and lilial (lily aldehyde) (Rodriguez et al., 2015; Zeng et al., 2018).

We purchased a total of 60 products from May-December 2025 from a range of suppliers, including pharmacies and health stores (e. g. Chemist Warehouse, Healthylife, iHerb), supermarkets (e. g. Coles), general online marketplaces (e. g. eBay, Etsy, Temu, Amazon) and independent suppliers. Details of each product are provided in Table S1. We tested 55 repellents, including 43 topical repellents, four wristbands, three clothing stickers, three ultrasonic devices, one garment (permethrin-treated socks) and one repellent applied to clothing. The topical repellents contained 18 different primary active ingredients (defined as the first active ingredient listed) with at least three products chosen for each of the following primary active ingredients: DEET, picaridin, oil of lemon eucalyptus, IR3535 and citronella. Twenty-five of the products contained multiple active ingredients (including synergists), though some products lacked a full ingredient list, did not specify the active ingredient, or made no distinction between active and inert ingredients that could also contribute to repellency (e.g. fragrances). Five products were identified as multi-purpose, with the mosquito repellent also labelled as a sunscreen, perfume and/or antiseptic cream. Forty-five of the products were explicitly labelled as a mosquito or insect repellent while the remaining ten were more ambiguously labelled (e.g. “keeps bugs away”, “reduces mosquito appeal”, “anti-bug”, “mozzie spray”), though all of these were advertised as being a mosquito repellent on the product website. Nearly half (25/52) of the non-exempt repellents tested were not registered with the APVMA, with all of these purchased online.

We also tested five products that were not labelled as mosquito repellents. We included Victoria’s Secret Bombshell which is reported to contain lilial and hedione, both of which show mosquito repellent efficacy (Rodriguez et al., 2015; Zeng et al., 2018). We also tested Calvin Klein CK One which contains a high concentration of hedione (Schaefer, 2015). We note that lilial was banned from use in cosmetics in the European Union in 2022 (Vieira et al., 2024) and may no longer be used in the formulation of Victoria’s Secret Bombshell tested in our study. Finally, we tested a small range of deodorants and antiperspirants based on previous evidence that common compounds in deodorants can affect mosquito attraction (Verhulst et al., 2016).

### Mosquitoes

We used wild-type *Ae. aegypti* mosquitoes in all experiments. We chose *Ae. aegypti* over other human-biting mosquito species that are more widespread in Australia (e. g. *Ae. notoscriptus*) due to *Ae. aegypti* being strongly anthropophilic and relatively easy to produce in large numbers for experiments. Populations were sourced from several suburbs throughout Cairns, Queensland, Australia in February 2024 and were at generations 18-22 in the laboratory at the time of the study. To remove *Wolbachia* endosymbionts present in the population (Hoffmann et al., 2011), adults were treated with tetracycline hydrochloride (2mg/mL) in 10% sucrose solution for the first two generations following laboratory establishment, returning the population to its originally *Wolbachia*-free state. The removal of *Wolbachia* was verified through qPCR assays (Lee et al., 2012).

*Aedes aegypti* mosquitoes were maintained in the laboratory at 26°C with a 12:12 light:dark cycle according to protocols described previously (Ross et al., 2017). Female mosquitoes were blood fed on a single human volunteer to obtain eggs for experiments. Blood feeding of *Ae. aegypti* on human volunteers was approved by the University of Melbourne Human Ethics committee (project ID 28583) and informed consent was obtained from the subject. Female *Ae. aegypti* used for arm-in-cage repellent trials were 5-25 days old and were provided with 10% sucrose solution and water prior to assays.

### Arm-in-cage trials

Trials ran from September 2025-August 2026 across eight adult human subjects (three male, five female). Each human subject tested all 60 products in a random order, with each product tested once per subject, for a total of 480 trials. To capture natural variation between human subjects, subjects were not instructed to alter their daily routines (e. g. diet, hygiene) or refrain from using deodorant, except for products applied to the forearm.

To test each product, we used a standardized arm-in-cage assay adapted from WHO guidelines (Organization, 2009), with modifications depending on the product category tested (Figure 1). Trials were conducted indoors at room temperature (∼ 22°C, 45% relative humidity). Human subjects tested each product on a separate day with a separate batch of mosquitoes, except for products that did not provide complete initial protection. In cases where multiple products were tested on the same day, subjects washed their arms thoroughly with soap and tap water and waited for at least 1 hr before testing a different product. Control and treatment arms were alternated between left and right for each product tested by the subject. Nitrile gloves were worn in all trials to ensure that only forearms were exposed to mosquitoes.

**Figure 1.**
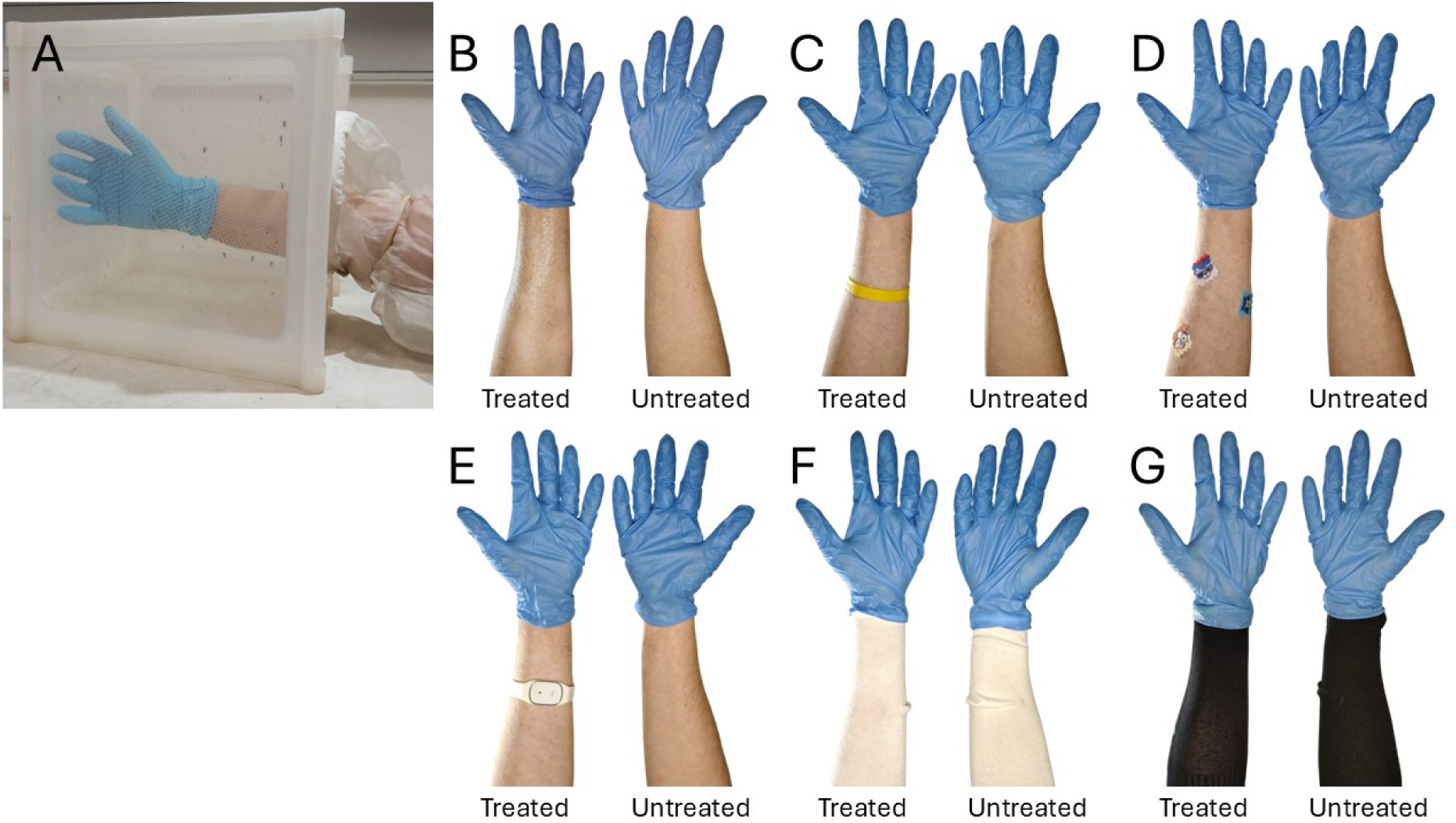
Arm-in-cage assays for testing different mosquito repellent product categories, showing treated and untreated control arms. (A) Arm-in-cage assay showing a treated forearm introduced into a cage of 50 *Ae. aegypti* mosquitoes. (B-F) Examples of treated and untreated forearms for testing different product categories: (B) topical repellents and perfumes, (C) wristbands, (D) stickers, (E) ultrasonic devices, (F) fabric repellents, (G) permethrin-treated socks. Comparisons in B-F involve landing assays while comparisons in G involve a blood feeding assay.

For each trial, repellents were applied immediately before introducing the treated forearm into a 30 x 30 x 30 cm cage (BugDorm-1, Australian Entomological Supplies, NSW, Australia) containing 50 female *Ae. aegypti* (Figure 1A). The number of landings (where mosquitoes landed on the skin for more than 3 seconds) was recorded for 3 min. Mosquitoes were not allowed to ingest blood, thereby minimizing exposure to mosquito bites. Before each test, an untreated control arm was introduced into the cage, and the number of landings was recorded in the same fashion as the treated arm. If fewer than 30 mosquitoes landed on the control arm in any test (indicating that mosquitoes were inactive), trials were repeated with a different batch of mosquitoes to ensure that estimates of repellency were reliable. To evaluate protection across time, this process was repeated every hour until more than 5 mosquitoes landed on the treated arm in two consecutive tests, or until 8 hours elapsed.

Relative protection at each time point was calculated by subtracting the number of mosquitoes landing on the treated arm from the number landing on the control arm, then dividing this value by the number of mosquitoes landing on the control arm. If more than 5 mosquitoes landed on the treated arm in the initial test, the test was repeated immediately and data from the two tests were averaged to calculate relative initial protection. Complete protection times are typically calculated by the time to the first mosquito landing (Organization, 2009). However, this approach is sensitive to differences in landing frequencies between tests and may also be prone to experimenter error. We therefore defined the time to loss of complete protection conservatively as the time until relative protection fell below 0.95. This threshold is equivalent to 5 mosquitoes landing on the treated arm compared to 100 on the control arm, or a 95% reduction in landing counts.

For topical repellents and perfumes (Figure 1B), human subjects were instructed to apply the product liberally to their forearm (from wrist to elbow) with the control arm left untreated. Topical repellents and perfumes were not reapplied during the trial. For wristbands (Figure 1C), products were placed around the centre of one forearm, with the control arm left untreated. Stickers (Figure 1D) were applied to the forearm, with three stickers per treated arm and none for the control. Ultrasonic devices (Figure 1E) were worn on the centre of the forearm or placed at the bottom of the cage (for non-wearable devices). The same arm was used for both the treatment and control, but with the device turned on for the treatment and off for the control. For the repellent applied to clothing (Figure 1F), human subjects wore socks covering both forearms for the duration of the trial. The repellent was applied liberally to the sock on the treatment arm but not applied to the control arm. White socks were used to facilitate landing counts.

For the permethrin-treated socks (Figure 1G), we initially followed the procedure for the repellent applied to clothing. However, pilot experiments indicated that the garment did not prevent mosquitoes from landing but did reduce blood feeding. To provide a more meaningful indication of exposure to mosquito bites, we instead used a blood feeding assay. Two cages of approximately 50 female mosquitoes were set up per subject. Subjects wore the permethrin treated sock on one arm and an untreated sock on the other arm, with no bare skin exposed. One forearm was placed in each cage and mosquitoes were allowed to blood feed for 15 minutes, after which the number of blood fed and unfed mosquitoes in each cage were counted. Trials were completed at a single time point for each human subject. Protection was calculated by subtracting the proportion of mosquitoes that fed on the treated arm from the proportion that fed on the control arm, then dividing this value by the proportion of mosquitoes fed on the control arm. All human subjects involved in the repellent tests provided their informed consent to participate in blood feeding.

## Statistical analysis

All statistics were performed using R in RStudio version 2025.05.1 (Posit team, 2025). We first considered mosquito landing counts in control tests where repellents were not applied. Generalized linear models on log transformed numbers were performed using the stats package (Team et al., 2018) with the car package (Fox et al., 2012) for Type II ANOVAs to test the impact of time interval, product and human subject on landing counts in control tests. We then explored the effects of human subject on the distribution of control counts, considering standard deviations, skewness and kurtosis which were computed using the e1071 package (Meyer et al., 2019). We used bootstrapping to compute the 95% confidence intervals of each of these parameters using the boot package (Canty & Ripley, 2017).

We determined relative protection for each product at different time intervals. Trials were discontinued when 5 mosquitoes landed on the treated arm in two consecutive tests, resulting in fewer comparisons across products at later time points. Almost all estimates of initial relative protection (proportion repelled relative to control) were between 0 and 1, however a small number of comparisons (15/480) had values <0, indicating that more mosquitoes landed on the treated arm than the control arm.

We first analysed each product separately to assess whether the product reduced mosquito landings compared with untreated control arms at the start of the trial (0 hr). Data were analysed using a two-stage generalized linear mixed model approach using the glmmTMB package (Magnusson et al., 2017). For each product, a binomial mixed model was used to test the probability of a mosquito landing on the treated or control arms, with subject and treatment included in the model. This was followed by a negative binomial mixed model for treatments where there were landings, with the same variables included to compare landing counts among observations. Effects of a product were estimated using estimated marginal means with the emmeans package (Lenth, 2018), comparing treated and control arms within each product. P-values were adjusted for multiple comparisons using the Benjamini– Hochberg false discovery rate correction.

We then used linear mixed effects models fitted with the lmerTest (Kuznetsova et al., 2017) and lme4 (Bates et al., 2015) packages to assess broader patterns of repellent efficacy across products in terms of loss of protection time (i. e. the time when landing counts on the treated arm were >5% of those on the control arm). Model assumptions were evaluated using residual-versus-fitted plots and normal Q-Q plots. We ran an initial analysis including product category and human subject as fixed effects with product included as a nested effect within categories, i. e. Loss of protection = category + product within category + human subject + interaction + error.

We then used a linear mixed effects model restricted to topical repellents to test for effects of topical repellent type, active ingredient and human subject. We also ran an analysis with APVMA registration as the fixed category effect due to some product categories being exempt from APVMA registration. We used Spearman correlations to examine the association between average complete protection time and advertised protection time where available.

Human subject effects were further investigated by computing Spearman correlations for complete protection time and relative efficacy of products at three time intervals (0, 1 and 2 hrs) between all pairs of human subjects. Similar rankings for the repellents between subjects would be expected to produce strong positive correlations. We also tested if mean landing counts on the control arm and mean complete protection time were correlated across human subjects by computing a Spearman correlation. Analyses for human subject effects were initially performed with perfumes excluded, though their inclusion had limited effects on the outcomes of all analyses and so these were retained to maintain consistency with the other analyses.

## Results

### Mosquito landings in untreated controls

We first assessed mosquito landing counts in the control arms where repellents were not applied by fitting a generalized linear model including product, time interval, and human subject as fixed effects. Across all tests we observed substantial variation in mosquito landing counts, ranging from 30-361 within 3 min (trials where fewer than 30 landings occurred in any test were discarded and repeated). Product (χ² = 341.59, df = 58, P < 0.001) and human subject (χ² = 413.66, df = 7, P < 0.001) had strong effects on control counts, whereas time interval (χ² = 19.13, df = 8, P = 0.014) had a comparatively small but significant effect, though landing counts did not show any clear increase or decrease across time (Figure S1). The distributions of control landing counts also differed markedly between human subjects (Figure S2), with significant differences in terms of their standard deviation, skewness and kurtosis based on non-overlapping bootstrap 95% CIs (Figure S3). This variation in landing counts and the presence of occasional mosquito landings on treated arms for products with high efficacy (Figure 2) led us to define the time to loss of complete protection as the point where the reduction in mosquito landings in the treated arm fell below 95%, rather than by the time to the first mosquito landing.

**Figure 2.**
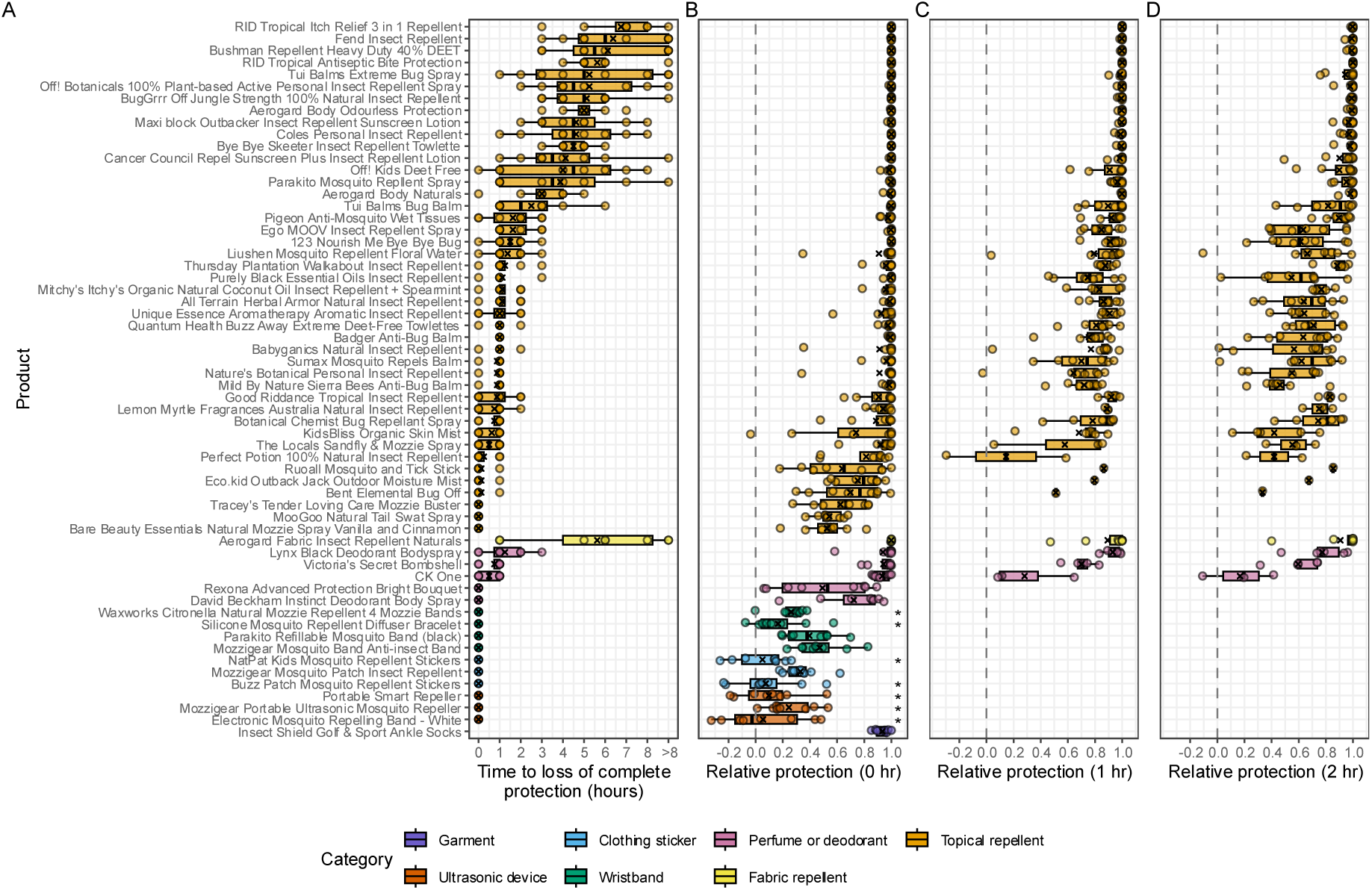
Efficacy of mosquito repellent products and perfumes against *Aedes aegypti* mosquitoes. (A) Time to loss of complete protection, defined as the first time point where the relative number of landings in the treated arm compared to the control arm was >5%. (B-D) Protection relative to untreated control arms at (B) 0, (C) 1 or (D) 2 hours post-application, where a value of 1 indicates complete protection relative to controls, 0 indicates no protection (dashed vertical lines) and negative values indicate a greater number of landings on the treated arm compared to the control. Products that did not significantly reduce the number of mosquito landings relative to the untreated control arm at 0 hr according to a negative binomial mixed model (P ≥ 0.05) are indicated with asterisks in panel B. P-values were adjusted for multiple comparisons using the Benjamini–Hochberg false discovery rate correction. Trials where initial complete protection was not observed are not shown in panels C and D. Boxplots show medians and interquartile ranges, X symbols represent means, while dots show data from individual human subjects. Products are ordered by category then mean time to loss of complete protection.

### Efficacy of individual repellent products

We then determined whether the individual products provided protection against mosquito bites directly following application (Figure 2). For the 59 products where landing counts were assessed, 17 provided complete initial (at 0 hr) protection against mosquito bites in trials across all eight human subjects (Figure 2B). An additional 27 products provided complete initial protection in some but not all subjects. Seven products (two clothing stickers, two wristbands and all three ultrasonic devices) did not significantly reduce the number of mosquito landings relative to the untreated control arm according to a negative binomial mixed model (Figure 2B). The permethrin-treated socks, which involved a blood feeding assay at a single time point rather than a landing assay, provided strong protection against mosquito bites (Figure 2B), though a small proportion of females successfully fed through the treated sock.

For products that provided initial protection of >95%, we performed additional tests at 1 hr intervals to determine the time when >95% protection was lost (Figure 2C, 2D; Figure S4). Twenty-six products had a mean time to loss of protection of more than 1 hr, while 17 were effective for more than 2 hrs (Figure 2A). The longest lasting products were all topical or fabric repellents, with RID Tropical Itch Relief 3 in 1 Repellent (DEET), Fend Insect Repellent (IR3535) and Bushman Repellent Heavy Duty 40% DEET (DEET) having the highest complete protection times overall (Figure 2A).

### Effects of product category, active ingredient, APVMA registration and advertised protection time on time to loss of complete protection

We investigated patterns of complete protection time across product categories and other characteristics (Figure 3). In an initial analysis of all products where loss of complete protection time was calculated, we found significant effects of both product category (linear mixed-effects model: χ² = 18.45, df = 5, P = 0.002) and human subject (χ² = 92.57, df = 7, P < 0.001). Topical and fabric repellents were the most effective product categories (Figure 3A) while wristbands, clothing stickers and ultrasonic devices were ineffective, with no products providing complete protection. However, two wristbands and one clothing sticker provided modest initial protection, with significantly fewer mosquitoes landing on treated arms compared to control arms at 0 hr (Figure 2B). Most perfumes and deodorants provided strong initial protection that declined rapidly (Figure 2, Figure 3A). Lynx Black deodorant was the only non-repellent product that provided complete protection for longer than 1 hr on average (Figure 2A).

**Figure 3.**
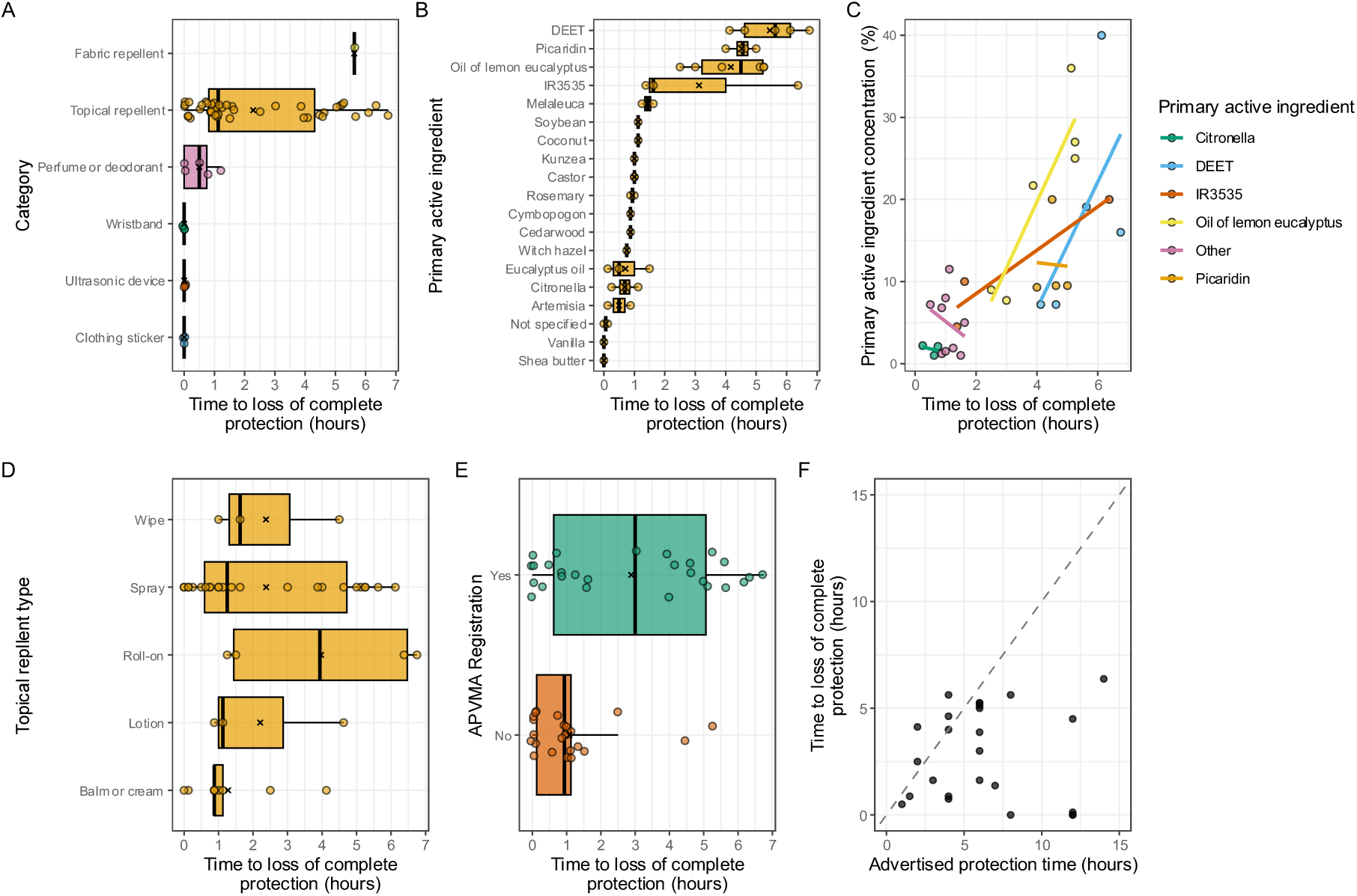
Time to loss of complete protection by (A) product category, (B) primary active ingredient, (C) concentration of active ingredient, (D) topical repellent type, (E) APVMA registration status and (F) advertised protection time. Boxplots show medians and interquartile ranges and X symbols represent means, with each data point representing the mean time to loss of complete protection across all human subjects, with one data point per product. Only topical repellents are included in panels B-D, while products exempt from APVMA registration (e.g. perfumes, ultrasonic devices) have been excluded from E. For panel F, the dashed line indicates parity between advertised and measured protection times. Two products with advertised protection times of 48 hr and complete protection times of 0 hr lie outside the axes and are not shown.

In an analysis that included only the topical repellents, we found a significant effect of primary active ingredient (linear mixed-effects model: χ² = 184.57, df = 18, P < 0.001). Topical repellents containing DEET, picaridin, oil of lemon eucalyptus and IR3535 as the primary active ingredient were the most effective overall (Figure 3B). Citronella and other essential oils were relatively ineffective as topical repellents, with only 6/25 of these products having mean times to loss of complete protection above 1 hr (Figure 3B) despite many providing strong initial protection (Figure 2B). We observed substantial variation among products of the same active ingredient (Figure 3B). This may partly reflect differences in the concentration of the active ingredients among products, with higher concentrations tending to have greater efficacy for several products where high concentrations were present (see regression lines in Figure 3C). Time to loss of complete protection was also affected by the type of topical repellent (χ² = 17.16, df = 4, P = 0.002), with roll-on products tending to perform better than other formulations (Figure 3D).

In a third analysis, we found a significant effect of APVMA registration on the time to loss of complete protection (linear mixed-effects model: χ² = 13.15, df = 1, P < 0.001), where registered products tended to be more effective overall (Figure 3E). However, we note that five registered products did not provide complete protection for any human subject (Figure 3E). For products that advertised a protection time, we found that only 5/28 met or exceeded their claimed efficacy (Figure 3F). There was also no clear relationship between advertised and measured efficacy (Spearman’s rho = –0.22, P = 0.252).

### Human subject effects on repellent efficacy

In earlier analyses we identified substantial variation in repellent efficacy between human subjects for the same product (Figure 2). Across all products, means for time to loss of complete protection (Figure 4A) and initial relative protection (Figure 4B) differed between subjects. Nevertheless, there were positive correlations between all human subjects in pairwise comparisons across the products for both complete protection time (Figure 4C) and initial relative protection (Figure 4D), suggesting that relative efficacy of the products were similar across subjects. Therefore, while overall efficacy differed between subjects, products that provided greater protection for one subject generally provided greater protection for other subjects, though this was not always the case (e.g. relative protection for Subject F at hr, Figure 4D). Positive pairwise correlations between human subjects were maintained at and 2 hr when considering products that provided complete initial protection (Figure S5).

**Figure 4.**
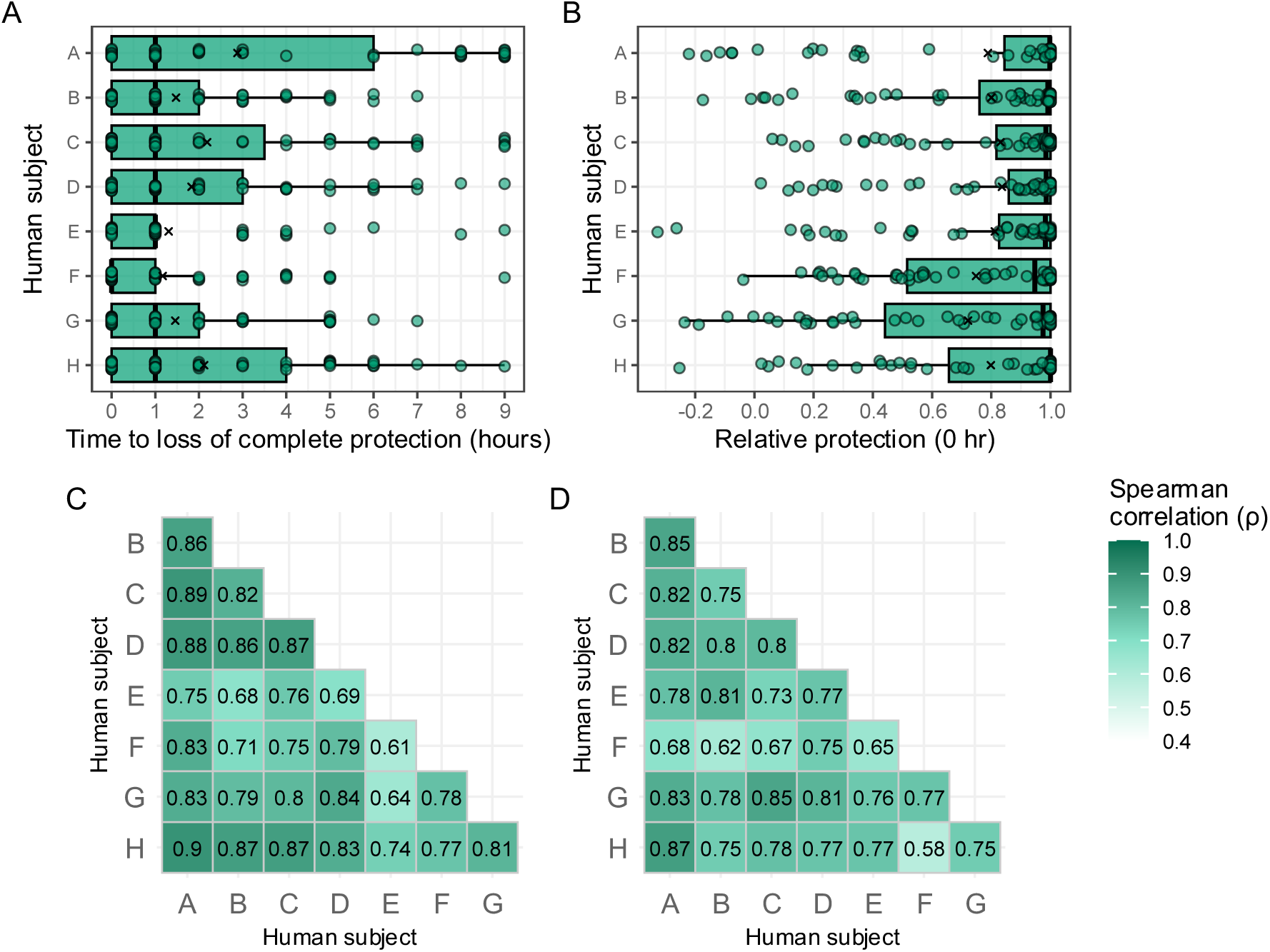
Human subject effects on mosquito repellent efficacy. (A) Time to loss of complete protection across all products, grouped by human subject. (B) Protection relative to untreated control arms at 0 hours post-application across all products, grouped by human subject. For A-B, boxplots show medians and interquartile ranges, X symbols represent means, while dots represent data from a single product. (C-D) Pairwise Spearman correlations between human subjects for (C) the time to loss of complete protection and (D) relative protection at 0 hr across all products. See Figure S5 for comparisons of relative protection at 1 and 2 hr between human subjects.

Perhaps surprisingly, we found a marginally significant positive correlation between mean landing counts on the control arm (Figure S1B) and mean time to loss of complete protection (Figure 4A) across the human subjects (Spearman: *ρ* = 0.74, P = 0.046, n = 8), suggesting that subjects with higher landing counts in the absence of repellents nevertheless showed longer complete protection times when compounds were applied.

## Discussion

Our comprehensive comparison of mosquito repellents on the Australian market demonstrates that many products are ineffective in preventing mosquito landings over a longer period, with only 17/54 providing more than two hours of complete protection in arm-in-cage assays. We also highlight the prevalence of unregistered products and many products that fail to meet their advertised protection times in our trials. Our comparisons reinforce the scientific consensus on which product categories and active ingredients are effective (Diaz, 2016; Faridah et al., 2026), with topical or fabric repellents containing DEET, picaridin, oil of lemon eucalyptus or IR3535 being the only products providing long-term complete protection, while wristbands, ultrasonic devices, clothing stickers and topical repellents containing essential oils are largely ineffective. We also demonstrate that some perfumes and deodorants can repel mosquitoes, consistent with previous studies (Rodriguez et al., 2015; Verhulst et al., 2016; Zeng et al., 2018), though we emphasize that complete protection was not long-lasting and that the perfumes were not applied according to their typical use.

Aside from clear differences in efficacy between product categories and active ingredients, we also found substantial variation among products with the same active ingredient. For instance, wristbands containing citronella and IR3535 were ineffective at providing complete protection but some topical repellents containing these active ingredients were effective. Within topical repellents, this variation can be partially explained by the concentration of the primary active ingredient, but differences may also reflect other factors. Many products contained multiple active ingredients and other ingredients including synergists, fragrances and sunscreens, confounding effects of primary active ingredient concentration. The physical properties of topical repellents also differed, ranging from thick balms to wipes and sprays that left little residue, which likely led to differences in the amount of repellent applied or their absorption into the skin. We did find some differences between types of topical repellents, though these comparisons are confounded by the active ingredient which tended to have larger effects.

Our study raises concerns regarding the availability of mosquito repellent products in Australia that are unregistered and/or ineffective. Although we did not find any unregistered products at major pharmacies and supermarkets, they were readily available online and through independent retailers. The sale of insect repellents without APVMA registration in Australia is unauthorized and products may be recalled for failing to meet safety regulations (https://www.apvma.gov.au/regulation/recalls) (Niven et al., 2020). Our results also highlight that registration with the APVMA does not guarantee that a product will be effective. For example, all registered mosquito bands and stickers failed to provide complete protection against mosquito landings in arm-in-cage assays, with some having no discernible effect at all. Ineffective products, particularly certified ones, may provide a false sense of security to users, leading to an increased risk of mosquito bites. The fact that most advertised protection times were higher than the complete protection times in our experiments also raises questions about how advertised values are determined and highlights the need for independent testing of products. Given the many different approaches used for testing mosquito repellents that can differ markedly in outcome (Luker, 2024), a standardized, transparent process may prove beneficial for both APVMA registration and for consumers.

We acknowledge several limitations to our study. First, the large number of products included led us to compromise on replication and reduce the frequency of testing within each trial. We prioritised between-subject replication to capture anticipated variability in repellent efficacy between humans but this meant that we were unable to assess within-subject variation in repellent efficacy. A recent study using a single DEET repellent found a similar level of variability in complete protection time both within and between human subjects (Moreno-Gómez, Monsonís-Güell, et al., 2025) but this may vary depending on the active ingredient. The use of 1-hour testing intervals was also a deliberate choice to account for the anticipated high variability in complete protection times, but this led to relatively coarse estimates of complete protection times for products that had provided only short-term protection. The completion of trials over the span of several months also introduces an additional level of noise to the data, including potential effects of mosquito cohort and age, which varied between trials. Although our measures of repellent efficacy were determined relative to paired untreated control arms, landing counts on these control arms were quite variable. Our design did also not fully capture the efficacy of repellents across time because we stopped trials after confirming that complete protection was lost. Most products with complete protection times above 0 but below 1 hr still provided a partial reduction in landings relative to the control arm at 2 hr. While complete protection times are a standard method of comparison (Organization, 2009), partial protection can still be useful for a consumer.

We also acknowledge the inherent limitations of arm-in-cage assays which do not capture the full range of mosquito host seeking behaviors under field conditions (Luker, 2024). Mosquitoes can be repelled through various modes of action depending on the active ingredient (Afify et al., 2019; Dagar & Ramakrishna, 2024; Syed & Leal, 2008), which may lead to differential efficacy when tested in various types of assay (Meier et al., 2025). For instance, the efficacy of repellents that mask body odors may have been underestimated in arm-in-cage assays where mosquitoes are already in close proximity to blood sources. Furthermore, the use of a single cohort of mosquitoes per trial leads to repeated exposure of mosquitoes to the repellent-an unlikely scenario under field conditions. Mosquitoes can experience decreased fitness following repeated exposure (Mulatier et al., 2018) and on the other hand may also become habituated (Lazzari et al., 2026; Stanczyk et al., 2013; Vinauger et al., 2014). Although we did not find a strong effect of time interval on control landing counts, the use of repeated exposures may contribute to variation in control landings between different products.

It is also worth emphasizing the clear differences in both repellent efficacy and distributions of landing counts in control arms across human subjects. Humans can vary in their attractiveness to mosquitoes (De Obaldia et al., 2022; Logan et al., 2008), but there are relatively few studies that consider human subject effects on repellent efficacy (Fradin & Day, 2002; Moreno-Gómez, Monsonís-Güell, et al., 2025; Rutledge & Gupta, 1999). Past work has identified some factors including sex (Drago et al., 2025) and menstrual cycle stages (Moreno-Gómez, Abril, et al., 2025). In our study, repellents tended to be more effective on subjects with higher landing counts in the control arms where repellents were not applied. Although landing counts should not be interpreted as a direct measure of attractiveness to mosquitoes, our data do suggest that higher levels of mosquito activity are not associated with reduced repellent efficacy. In future it would be interesting to further test links between attraction and repellent efficacy across a larger panel of human subjects.

In summary, our study provides a new and comprehensive comparison of mosquito repellent products sold in Australia. It not only highlights the prevalence of unregistered and ineffective mosquito repellents but also provides evidence supporting the effectiveness of repellents containing DEET, picaridin, oil of lemon eucalyptus and IR3535 in preventing mosquito landings. Overall, while our data show clear differences in efficacy between products, we emphasize that our results reflect differences under a defined set of laboratory conditions. Field trials remain crucial for demonstrating efficacy in a more realistic context (Frances et al., 2009) and it is also important to consider further testing against local mosquitoes given that the sensitivity of mosquitoes to repellents can vary between (Afify & Potter, 2020; Barnard & Xue, 2004; Curtis et al., 1987) and within (Deletre et al., 2019; Stanczyk et al., 2010) species.

## Acknowledgements

We thank members of the Pest and Environmental Adaptation Research Group for discussions on mosquito repellents and Xinyue Gu, Belinda van Heerwaarden and Anne Hillerman for their input in identifying products. We thank Scott Ritchie for providing the mosquito population used in this study.

## Supplementary information

**Table S1.** Details of products included in the study, including summary data on complete protection times and relative protection at 0, 1 and 2 hr averaged across human subjects.

**Table S2.** Repellent efficacy data for each combination of product and human subject. Data on time to loss of complete protection were calculated from the full dataset in Table S3.

**Table S3.** Repellent efficacy data for each individual test. Data are presented separately for each combination of product, human subject and time interval.

**Figure S1.**
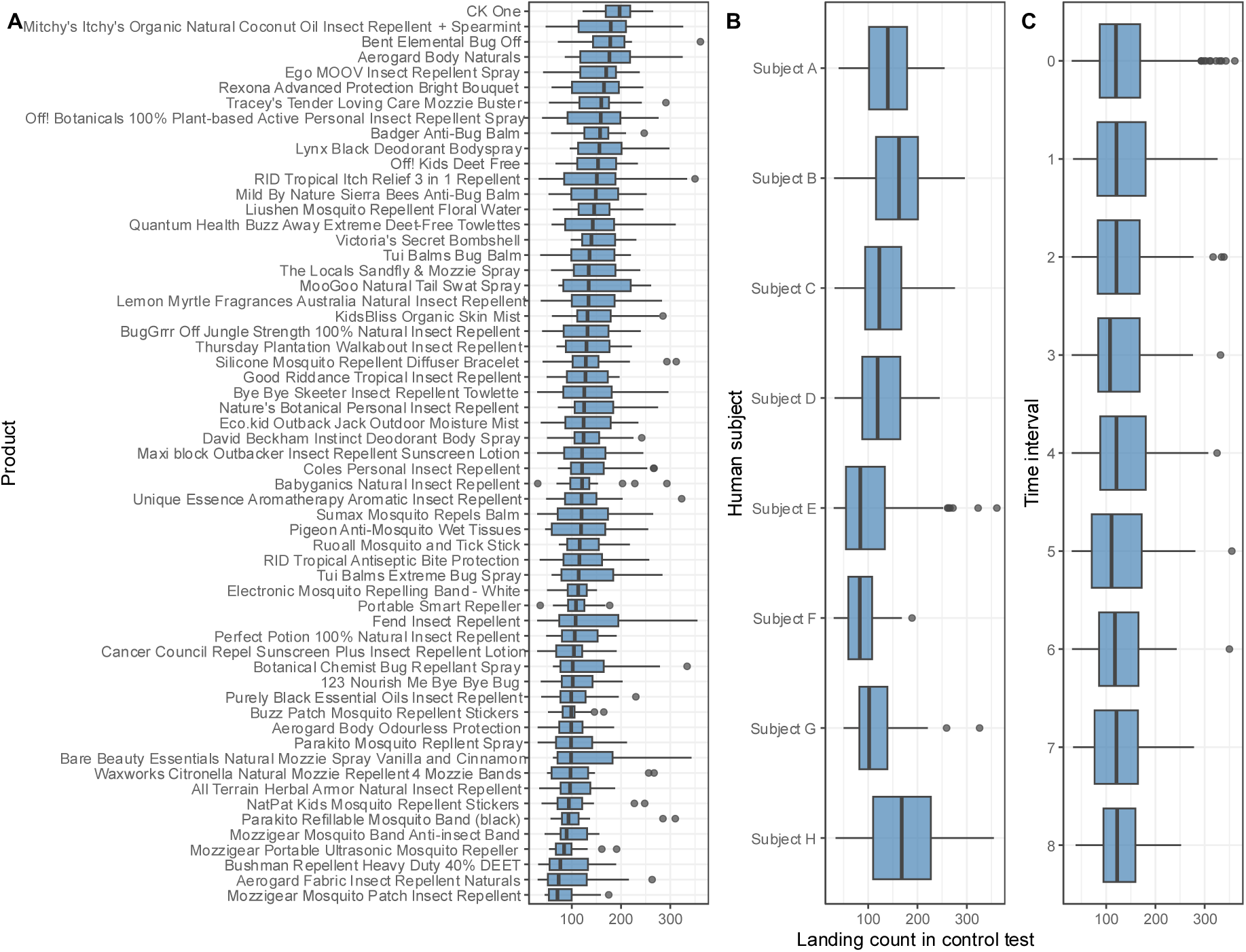
Mosquito landing counts in control tests for each (A) product, (B) human subject and (C) time interval. Boxplots show medians and interquartile ranges. Products in panel A have been ordered by median landing count.

**Figure S2.**
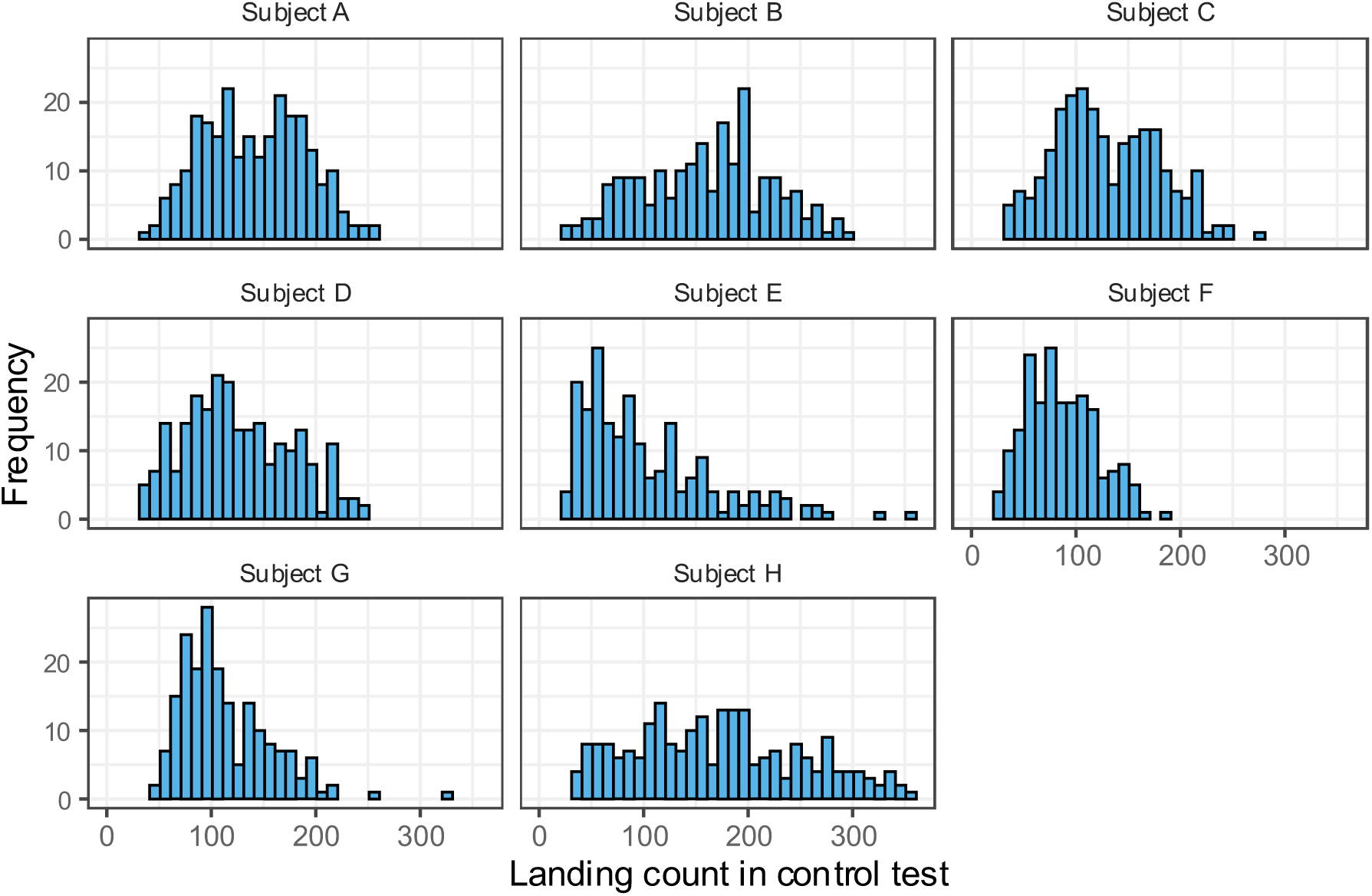
Histograms of mosquito landing counts in control tests where repellent products were not applied. Data are presented separately for each human subject.

**Figure S3.**
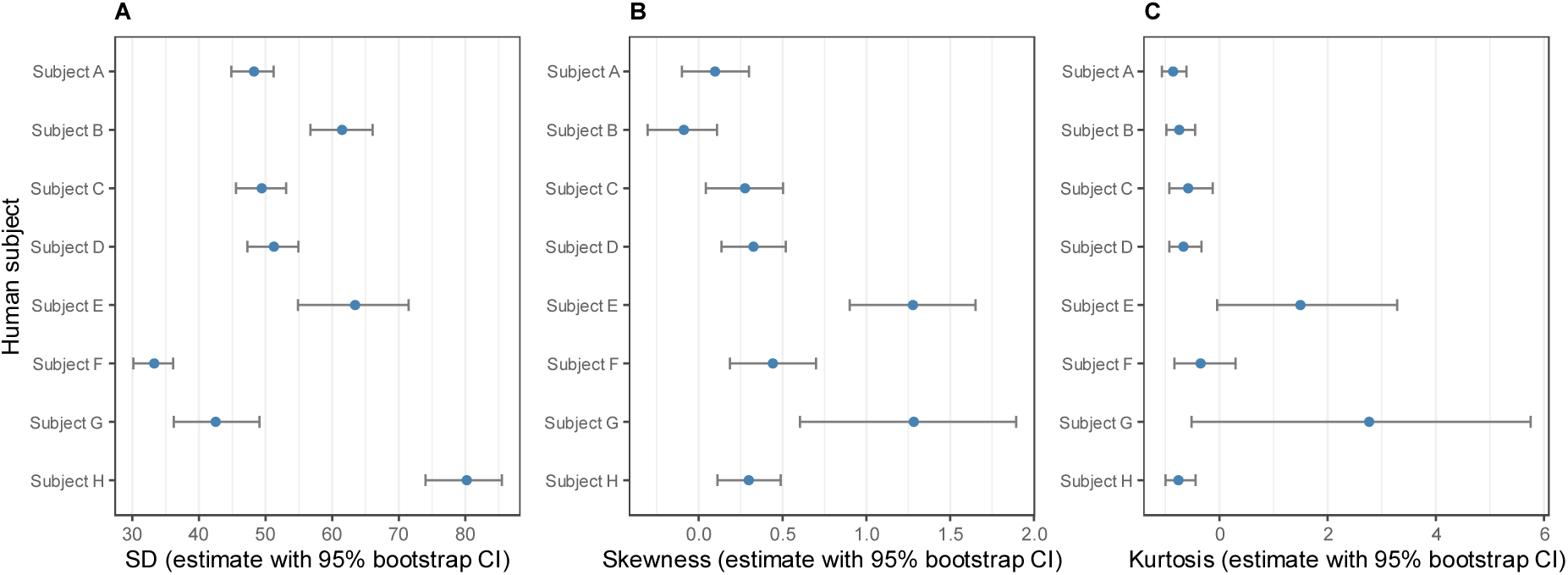
Descriptive statistics for mosquito landing counts in control tests across each human subject. Estimates and 95% bootstrap CIs are presented for (A) standard deviations, (B) skewness and (C) kurtosis.

**Figure S4.**
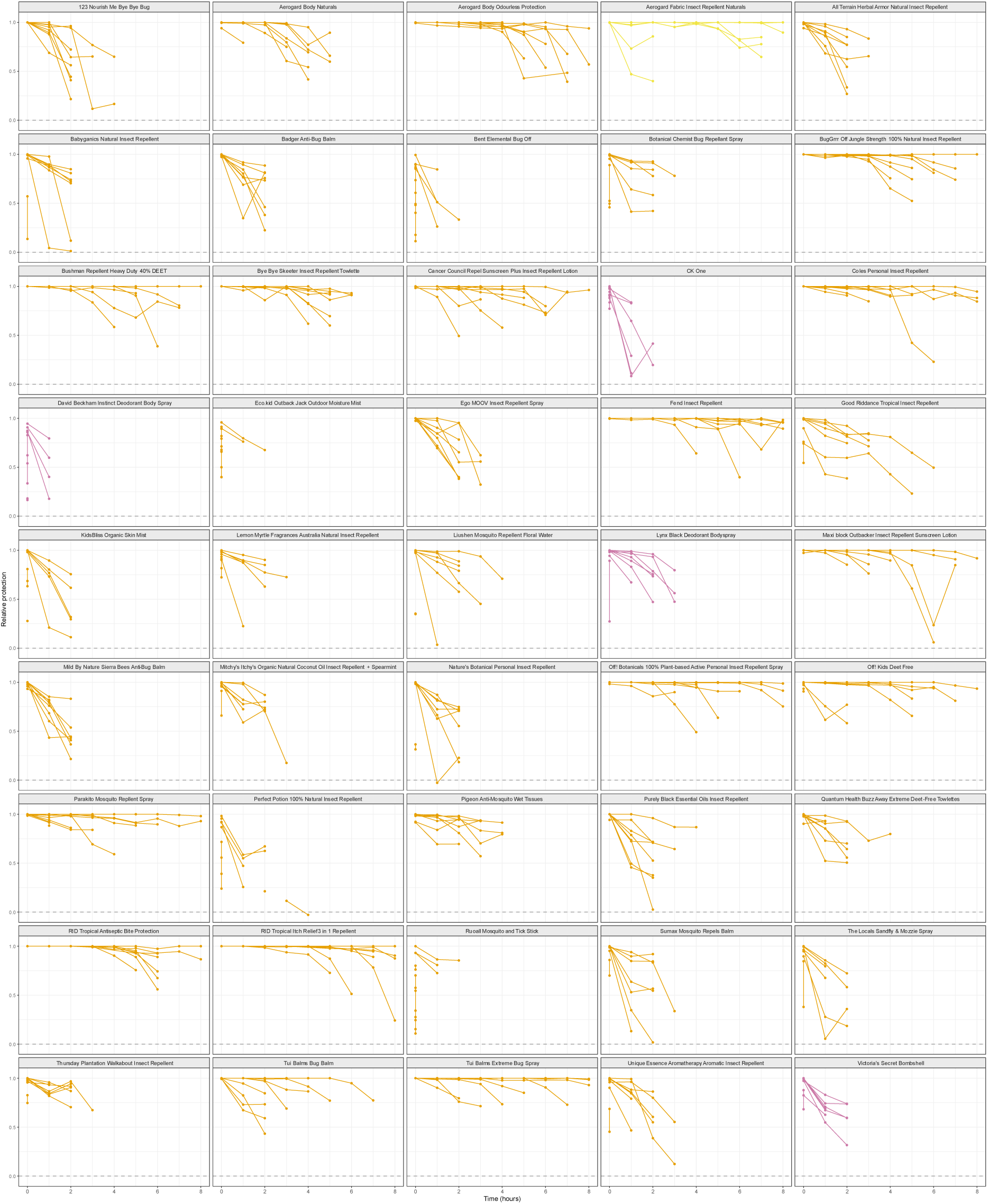
Efficacy of mosquito repellent products and perfumes across time. Each panel shows a different product that provided complete initial protection in at least one trial. Protection was calculated relative to untreated control arms at 1-hour intervals until 5 or more mosquitoes landed on the treated arm in two consecutive tests. Each line represents a single trial from a different human subject. A value of 1 indicates complete protection, 0 indicates no protection and negative values indicate a greater number of landings on the treated arm compared to the control.

**Figure S5.**
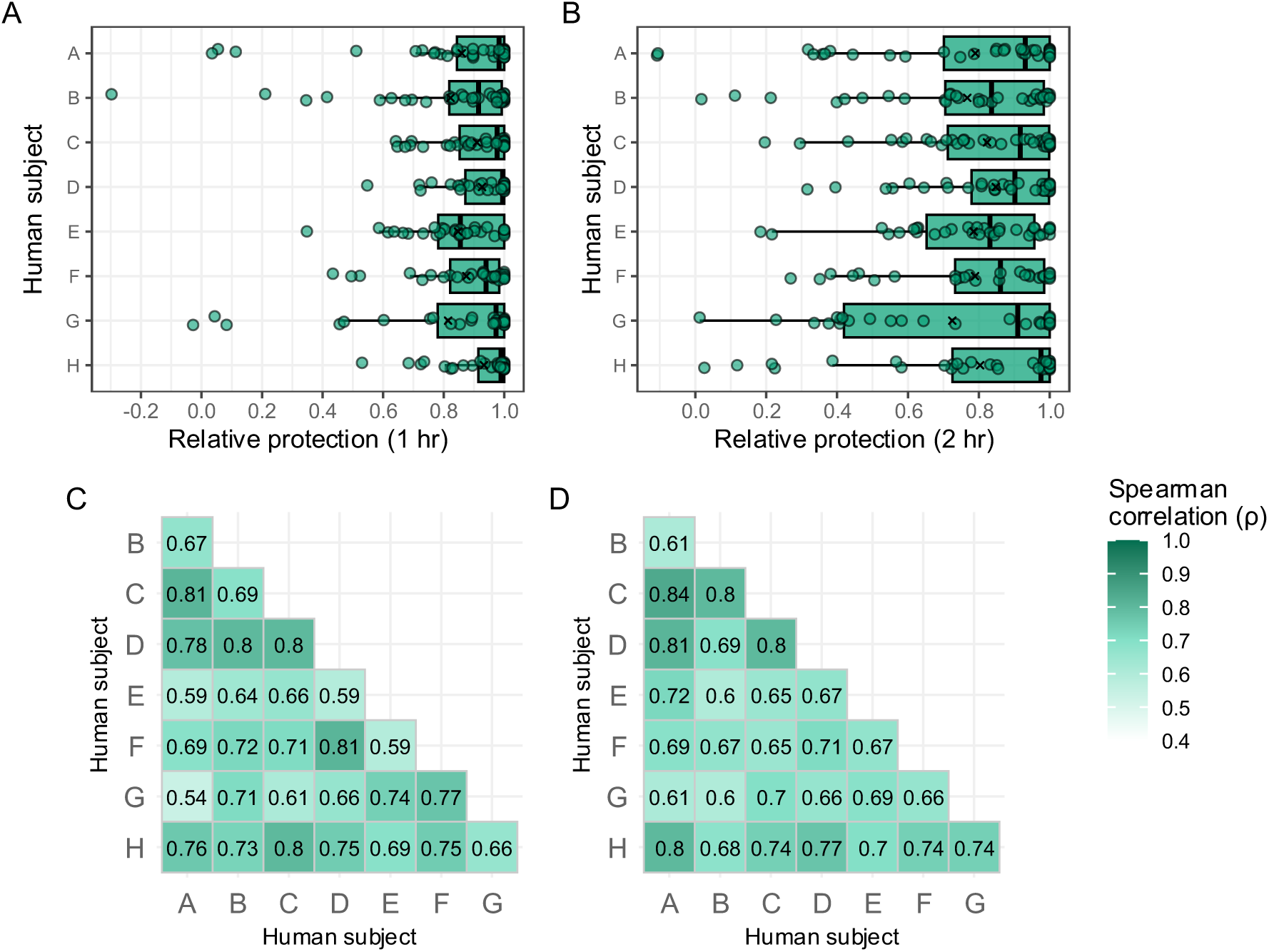
Human subject effects on mosquito repellent efficacy at 1 and 2 hrs post-application. (A-B) Protection relative to untreated control arms at (A) 1 hour and (B) 2 hours post-application across all products, grouped by human subject. For A-B, boxplots show medians and interquartile ranges, X symbols represent means, while dots represent data from a single product. (C-D) Pairwise spearman correlations between human subjects for relative protection at (C) 1 and (D) 2 hr. Comparisons where no landings occurred on treated arms were included.

